# Typical somatomotor physiology of the hand is preserved in a patient with an amputated arm

**DOI:** 10.1101/2021.02.12.430936

**Authors:** Max van den Boom, Kai J. Miller, Nicholas M. Gregg, Gabriela Ojeda, Kendall H. Lee, Thomas J. Richner, Nick F. Ramsey, Greg A. Worrell, Dora Hermes

## Abstract

Electrophysiological signals in the human motor system may change in different ways after deafferentation, with some studies emphasizing reorganization while others propose retained physiology. Understanding whether motor electrophysiology is retained over longer periods of time can be invaluable for patients with paralysis (e.g. ALS or brainstem stroke) when signals from sensorimotor areas may be used for communication or control over neural prosthetic devices. In addition, a maintained electrophysiology can potentially benefit the treatment of phantom limb pains through prolonged use of these signals in a brain-machine interface (BCI).

Here, we were presented with the unique opportunity to investigate the physiology of the sensorimotor cortex in a patient with an amputated arm using electrocorticographic (ECoG) measurements. While implanted with an ECoG grid for clinical evaluation of electrical stimulation for phantom limb pain, the patient performed attempted finger movements with the contralateral (lost) hand and executed finger movements with the ipsilateral (healthy) hand.

The electrophysiology of the sensorimotor cortex contralateral to the amputated hand remained very similar to that of hand movement in healthy people, with a spatially focused increase of high-frequency band (65-175Hz; HFB) power over the hand region and a distributed decrease in low-frequency band (15-28Hz; LFB) power. The representation of the three different fingers (thumb, index and little) remained intact and HFB patterns could be decoded using support vector learning at single-trial classification accuracies of >90%, based on the first 1-3s of the HFB response. These results demonstrate that hand representations are largely retained in the motor cortex. The intact physiological response of the amputated hand, the high distinguishability of the fingers and fast temporal peak are encouraging for neural prosthetic devices that target the sensorimotor cortex.

## Introduction

Deafferentation from the loss of a limb affects the inputs and outputs to and from sensorimotor areas in the brain. However, it is not yet clear what happens to the physiology of these cortical regions when a limb is amputated. Using electrocorticography (ECoG) measurements in humans, it has been well established that hand movements cause a spatially focal increase in high frequency amplitude in the sensorimotor cortex and a spatially distributed decrease in low frequency amplitude (Crone, Miglioretti, Gordon, and Lesser 1998; Crone, Miglioretti, Gordon, Sieracki, et al. 1998; Miller et al. 2007; Hermes, Miller, et al. 2012). Furthermore, the individual finger movements can be distinguished topographically using the high frequency signals (Miller et al. 2009; Siero et al. 2014). It is unknown whether these basic physiological changes are maintained after the amputation of a limb.

Several studies have reported that sensorimotor areas reorganize after amputation in humans using transcranial magnetic stimulation (Cohen et al. 1991; Röricht et al. 1999) and fMRI (Elbert et al. 1994; Dettmers et al. 2001; Lotze et al. 2001), and in macaque monkeys using electrical stimulation (Qi et al. 2000). These studies suggest that areas previously related to the amputated limb can associate with other muscle groups. However, another fMRI study on upper arm amputations shows that some form of representation is preserved, even over longer periods of time (Bruurmijn et al. 2017). Preserved motor physiology would be invaluable for specific clinical purposes such as Brain Computer Interfacing (BCI). Using BCIs, people with paralyses can use the electrophysiological signal from the brain, generated by attempted hand movement, to control communication devices (Vansteensel et al. 2016) or other assistive devices (Benabid et al. 2019). Furthermore, establishing BCI control could potentially help reduce phantom limb-pain (Yanagisawa et al. 2020), it is therefore important to understand the extent to which motor physiology is preserved.

In this study we were provided with a unique opportunity to investigate sensorimotor physiology with ECoG measurements in a patient with an amputated arm. The patient was implanted with an ECoG array for clinical evaluation of phantom limb pain and we measured ECoG signals during attempted finger movements of the contralateral, lost, hand. We found that the typical spatio-temporal organization of hand-movement physiology was preserved and that information of separate finger representations was retained.

## Methods

### Participant

A 62-year-old male with a left above-elbow amputation secondary to a snowmobile accident underwent temporary placement of a subdural electrode array for a trial treatment of phantom limb pain by electrical subdural cortical stimulation (Krushelnytskyy et al. 2019). Experimental data were collected during breaks in trials of different electrical stimulation parameters over a period of 10 days. The patient was right-handed and had his left-arm amputated 3 years and 11 months before ECoG grid implantation. The patient reported waking from his initial left mid-forearm amputation with phantom arm and hand pain, and that his pain has persisted since that time. He underwent two additional surgeries, ultimately completing a shoulder disarticulation and full humeral amputation. His pain has resulted in functional impairment and reduced quality of life despite trials of opiate medications, mirror therapy, an intensive pain rehabilitation program, and treatment with an implanted spinal cord stimulator. After the monitoring period, the patient was equipped with a cortical stimulator. At a six months follow up, he reported that the phantom limb pain dropped from 8-9/10 severity to a typical range of 5-6/10, and no side effects were reported.

The study was approved by the Institutional Review Board of Mayo Clinic (IRB 15-006530) and the patient provided informed consent to participate in the study, in accordance with the declaration of Helsinki (2013).

### Recordings

An electrode array of 36 circular platinum contacts (AdTech, 6 x 6 electrodes, 2.3 mm exposed diameter, 10mm inter-electrode distance) embedded in a silastic sheet was surgically placed over the fronto-parietal region, including the sensorimotor cortex (Figure 1A). Electrodes were localized using a high-resolution CT-scan and projected (Hermes et al. 2010) onto a cortical surface rendering generated from the preoperative anatomical T1 weighted MRI scan (GE 3T Discovery). During recording, all electrodes were referenced to an inactive subgaleal electrode with the recording surface facing away from the brain. The signals were amplified and digitalized at 2048 Hz. Upon inspection of the electrode signals, two channels that contained severe noise were excluded from analysis.

**Figure 1.**
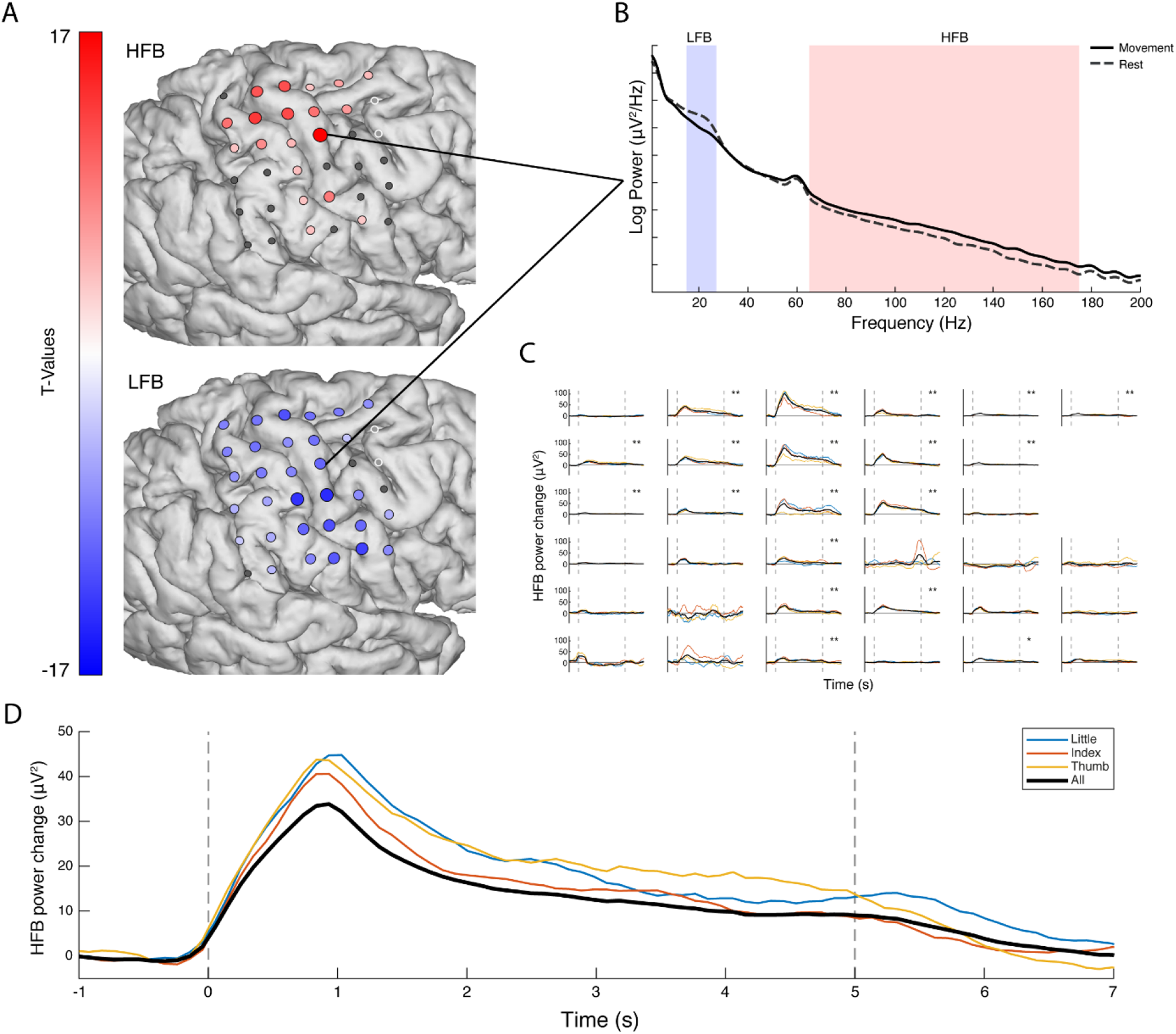
The electrophysiological response during attempted movement versus rest. **(A)** The changes in HFB (top, 65-175Hz) and LFB (bottom, 15-28Hz) for each grid electrode. Electrodes with a red or blue color had a significant change in band-power, whereas electrodes with in insignificant change in band-power are shown in grey; the two excluded electrodes are shown in white. **(B)** The power spectra of movement (solid line) and rest (dashed line) for a single electrode. **(C)** The HFB power changes over time, each graph represents one electrode. The black line represents all fingers, whereas the colored lines represent individual fingers. The two vertical dotted lines indicated the cue on- and offset. **(D)** The HFB power changes over time were averaged across those electrodes that showed a significant increase for a condition (blue: little, red: index, yellow: thumb, black: all fingers). The black trace has a lower amplitude because a different set of significant electrodes contributed to each trace, with more significant, but lower amplitude electrodes contributing to the black trace.

### Tasks

The subject was presented with two tasks: an attempted and executed movement task. During the attempted movement task, the subject was asked to attempt finger movements with the (left, contralateral) amputated hand; During the executed movement task the subject was asked to perform finger movement with the (right, ipsilateral) hand on the healthy arm. Both tasks featured the exact same design with 5s of finger movement and 3s of rest. The subject was cued via a bedside monitor with a picture of a hand and asked to (attempt to) move one of three fingers: the thumb, index or little finger. Each run of a task featured 15 movement cues for each finger, resulting in 45 randomized trials per run. The subject performed two runs of attempted movement and two runs of executed movement.

### Analysis

The analysis and classification routines were implemented using custom MATLAB (Mathworks inc.) code that is provided alongside this article at: https://osf.io/vmxdn/. Before analysis, the two runs of each task were concatenated and a small number of trials, that showed large artifacts in the signal or in which the patient was distracted, were excluded (2 trials were excluded for attempted movement and 4 trials were excluded for executed movement). The data were re-referenced to the common average by regressing the common average out from each channel.

### Spectral power change

The contralateral power changes during attempted hand movement were investigated by extracting an epoch of 1-4s after cue onset as movement. During executed movement (ipsilateral), the movement epoch was set to start 100ms before the actual movement of the healthy hand to the end of the actual hand movement based on the concurrent video; 100ms was subtracted to account for the delay between the cortical signal and initiation of the movement (Evarts 1973; Cheney and Fetz 1984; Miller et al. 2009). An epoch of 2s before cue onset was considered as rest in both attempted and executed movements. The power spectral density of each epoch was calculated every 1 Hz by Welch’s method (Welch 1967) with a 250ms window and an overlap of 125ms. A Hann window (Nuttall 1981) was applied to each epoch to attenuate the edge effects. Per channel, the resulting power spectra were log10 transformed and normalized to the mean power over all epochs at each frequency. The high frequency band (HFB) power was obtained by calculating the average power over 65Hz to 175Hz, whereas low frequency band (LFB) power was the average over 15Hz to 28Hz.

In order to plot the spectral power changes on the rendered brain surface, we calculated the T-statistics for both the HBF and the LFB per channel by testing the power of movement trials against the rest trials. A Bonferroni correction was applied while testing the T-values for significance.

To visualize the electrophysiological response over time, we filtered each electrode signal using a third-order Butterworth filter for either the high frequencies (HFB, 65-175Hz), or the low frequencies (LFB, 15-28Hz). After filtering, the power of the amplitude was calculated using a Hilbert transformation. Trial-epochs, of 2s before to 7s after cue onset, were extracted and normalized by subtracting the signal mean power of the 2s before cue onset from each individual trial. An average across each condition was calculated and smoothed with a moving average window of 1s.

### Temporal window for finger movement classification

In order to investigate to what degree spatial finger-representations were preserved and distinguishable, single-trial classification was performed on the individual fingers. For executed hand movement, we used the signal when the patient was moving the finger on the healthy hand. However, during attempted finger-movement there is no external behavioral measurement available to assess when (after cue onset) the patient started to attempt the movements, where in time the strongest decodable response occurs and whether such a response is transient or sustained. In order to address these factors, we split the data in two halves. One half was used to explore decodability of the response over time and optimize the time-window parameters for decoding. These time-window parameters were then used in the other half for further decoding analysis.

We explored the response over time and optimized it for decoding by applying several different time-windows while decoding the finger-movement, thereby restricting the information available to the classifier. These time-windows differed in size from 250ms to 5000ms and in placement from cue onset, ranging from the beginning to the end of the trial.

An optimal time-window for classification was determined by first applying gaussian smoothing to the classification accuracies over the window size (σ: 2.5) and offset (σ: .5) dimensions. Smoothing prevents the selection of parameters with local classification accuracy peaks in the parameter optimization half of the data, and allows for the selection of temporal parameters that are optimal in general. After smoothing, the time-window with the highest classification score was selected, and its offset and size were used for further classification analyses.

### Classification

For classification, the HFB power of the different channels at the trial-epochs were used as features in a Support Vector Machine (SVM) with a linear kernel (Bishop 2006). The HFB power was calculated per epoch in the same way as described above (i.e. using Welch’s method, log10 transformed and averaged over 65-175Hz), except that the spectra were not normalized. The unnormalized HFB power was used since the SVM maps each input feature to its own (scaled) dimension, and allows us to classify on the movement-epochs alone. We achieved multi-class classification (3 fingers) by applying a one-versus-all classification scheme in which every class is classified against the data of the other classes together and the winner (that is furthest from the hyperplane) takes all. A leave-one-out cross validation was used and resulted in a classification accuracy score, which is the percentage of trials predicted correctly. Classification scores were empirically tested for significance using a Monte Carlo distribution based on 100.000 permutations (Combrisson and Jerbi 2015).

### Spatial analyses

Searchlight analyses were performed to establish which area on the cortex was most informative for attempted finger movement and how many electrodes (i.e. which grid configurations) would be needed to reliably classify the individual fingers.

The most informative cortical regions to decode attempted finger movements were identified using a random search procedure. During the random search procedure, a subset of 1 to 36 electrodes was selected at random to classify from. This procedure was repeated 10.000 times and, for each electrode, the average accuracy over all iterations was calculated and z-scored.

Searchlight analyses were performed to identify which anatomical scale of coverage would provide the most information for classification. During the searchlight analyses, a searchlight (i.e. a block of electrodes) was used for classification. The searchlight, with a fixed block size (e.g. 2×2 electrodes) was placed at every possible position within the grid. Afterwards, for each electrode, the average over all the iterations in which that particular electrode was involved was calculated. Searchlight analyses were performed with all possible searchlight sizes and shapes, representing grids of all different sizes (1×6/6×1 to 6×6 electrodes).

## Results

A typical electrophysiological motor response occurred upon attempted movement. In order to investigate to what extent the sensorimotor cortex showed typical physiology after deafferentation, we measured ECoG responses in a patient with an amputated arm. The patient reported vivid movements of the amputated hand and could describe in clear fashion how well the different fingers moved during the task. Figures 1A and 1B show the electrophysiological differences between attempted movement of the missing hand and rest. During attempted movement, a spatially distributed decrease of LFB power occurred. Simultaneously, significant focal increases of HFB power were found, most notably around the primary sensorimotor hand-areas. Strong decreases in power were observed in a narrow range of the lower frequencies (β band, 15-28Hz), for completeness, supplementary Figure 1 also illustrates the responses in the alpha range (8-13Hz). High frequency power increases were distributed over a broad range of higher frequencies (>65Hz) (Figure 1B).

During attempted movement there was no behavioral measurement available to determine when - after cue onset - the patient actually starts attempting the movement, nor where in time to expect the physiological response. Figures 1C and 1D present the changes in HFB power over time for each of the individual electrodes and the power averaged over all significant electrodes for each condition. A clear peak in HFB power is visible around 1s second after the cue onset for all fingers.

### Differential electrophysiological responses for the three individual fingers

Although each of the individual fingers provided a spatially different electrophysiological response, as shown in Figure 2, it is difficult to distinguish a clear topographical order.

**Figure 2.**
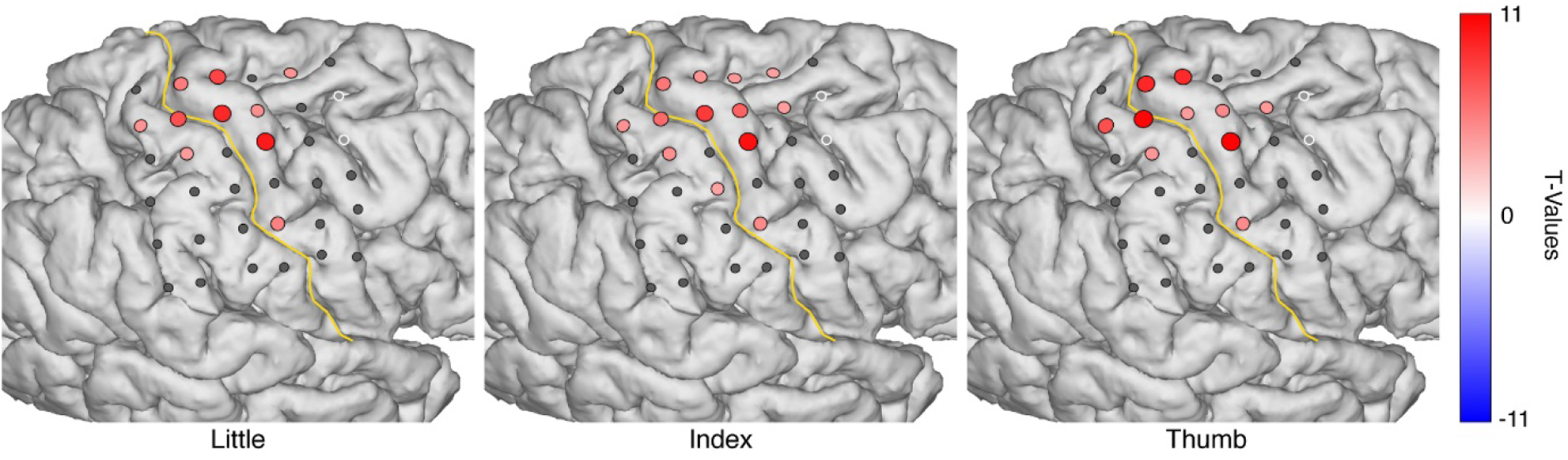
The HFB power changes upon attempted movement of each individual finger. Electrodes with a significant change in HFB band-power are shown in red, whereas electrodes with an insignificant HFB change are shown in grey, the two electrodes that were excluded are shown in white. The yellow line represents the central sulcus.

To investigate whether the representations of the individual fingers were preserved and sufficiently distinguishable we wanted to perform single-trial classification of the individual fingers. However, since there was no behavioral movement information available, we first needed to establish where in time the most information on attempted finger movement was present. By restricting the classification to the information within specific time-windows we could explore where in time the most information about attempted finger movement resided. Figure 3 shows the classification accuracies based on the HFB power within a specific window of time, ranging from 250ms to 5000ms, placing the window at different moments between the cue onset (t = 0ms) and offset (t = 5000ms).

**Figure 3.**
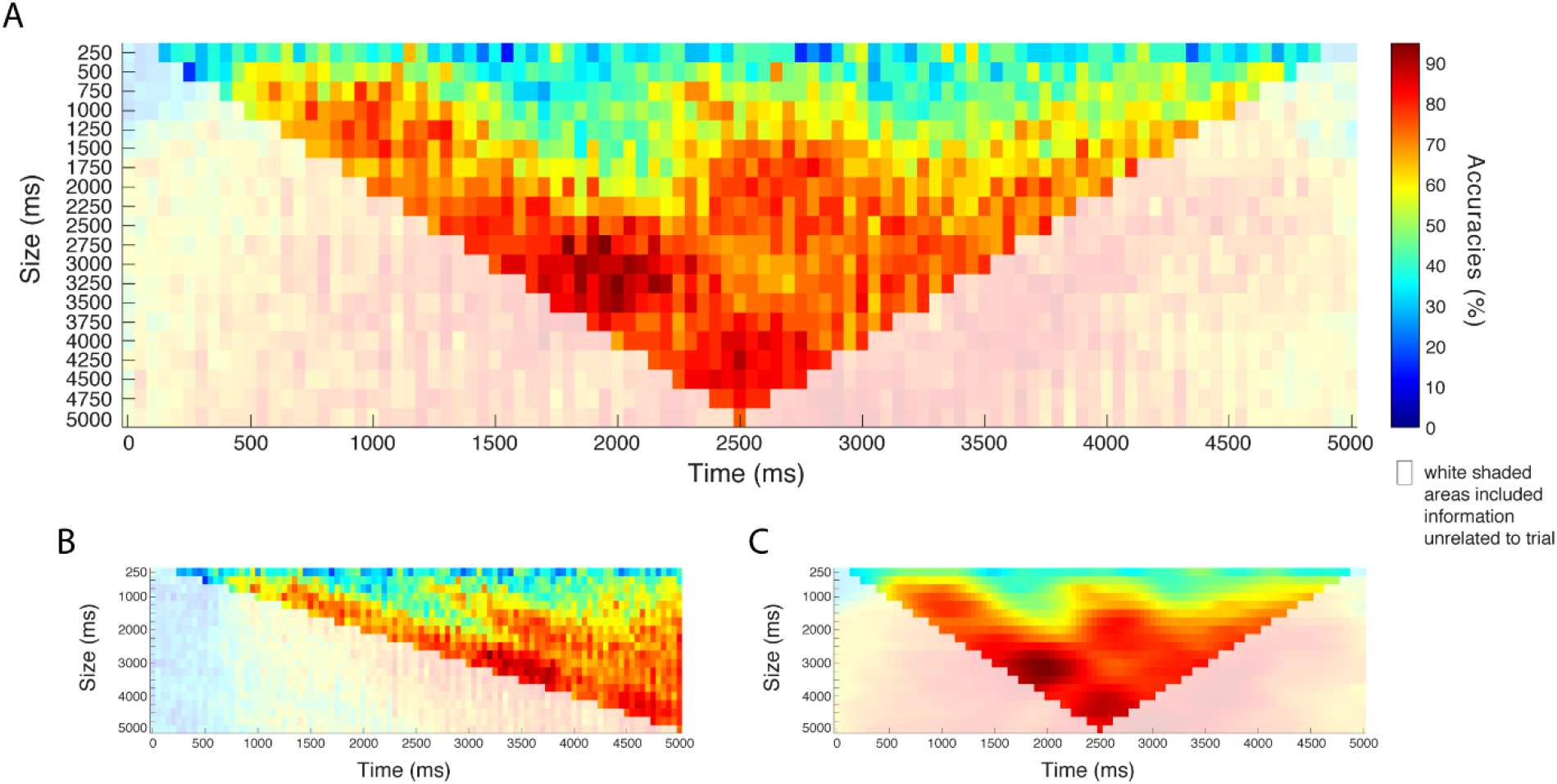
The classification accuracies at different window sizes (y-axes) and offsets from cue onset (x-axes), based on half of the data (~45 trials). **(A)** The x-axis indicates the center of each window. **(B)** The same plot shown, but with the x-axis indicating the right of each window, such that each time point includes the information present before that time. **(C)** Shows the classification accuracies smoothed with a Gaussian filter (offset σ: .5, size σ: 2.5). The white-shaded regions in each graph indicate the classification accuracies in which the window included information unrelated to the trial (i.e. rest before or after the trial).

Smaller time windows (<750ms) seem to provide less good decoding accuracies (0% - 50%) in comparison to medium (750 - 2500ms) or larger time-windows. Medium-sized windows can perform reasonably well (60%-80%) depending on their offset in time. Window sizes of about 2500ms to 5000ms performed well overall (> ~70%). In terms of window offset, the highest classifying windows take information from the beginning of the trial, regardless of window size. After Gaussian smoothing (Figure 3C) of the classification results, the optimal window (i.e. the highest classification score) was found at a width of 3000ms at 1950ms (window center) after cue onset, which converts to an epoch window from 450ms to 3450ms after cue onset. The remainder of the results are based on this epoch and are performed on the half of the data that was not used for parameter optimization.

The second half of the data showed that the classification accuracy of attempted finger movements based on the spatial features of the HFB power was significant at 93% (above 45% chance level calculated with Monte-Carlo simulation). The sensitivity values for the individual fingers (Little: 87%, Index: 93% and Thumb: 100%), shown in Figure 4, were also significantly above chance for all three fingers.

**Figure 4.**
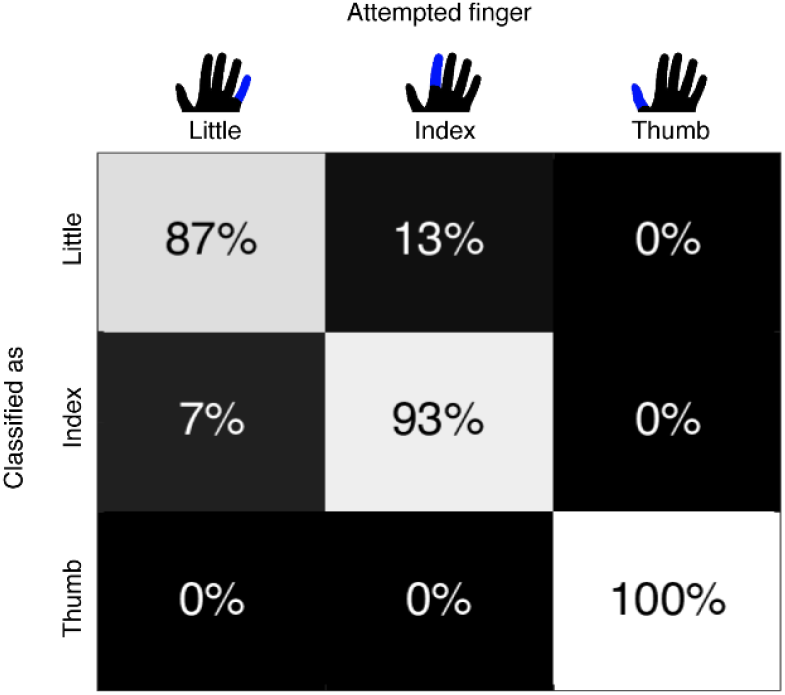
Confusion matrix with the classification scores of the individual fingers. Each column represents the finger of which movement was attempted. The rows represent how each of those finger movements was classified.

### Topographical organization of attempted hand movement information

For all of the finger movements we investigated where on the cortex the related activity was located. Figure 1 and 2 already showed that most of the HFB power changes related to attempted movement occur around the hand and arm region of the pre and post central gyrus. Significant HFB changes extended beyond pre and post central gyrus to more anterior premotor regions as well. The random search classification results in Figure 5C confirm that most information indeed resides in S1 and M1, specifically in areas of the pre and post central gyrus that are well known to represent hand and finger movements (Penfield and Boldrey 1937).

**Figure 5.**
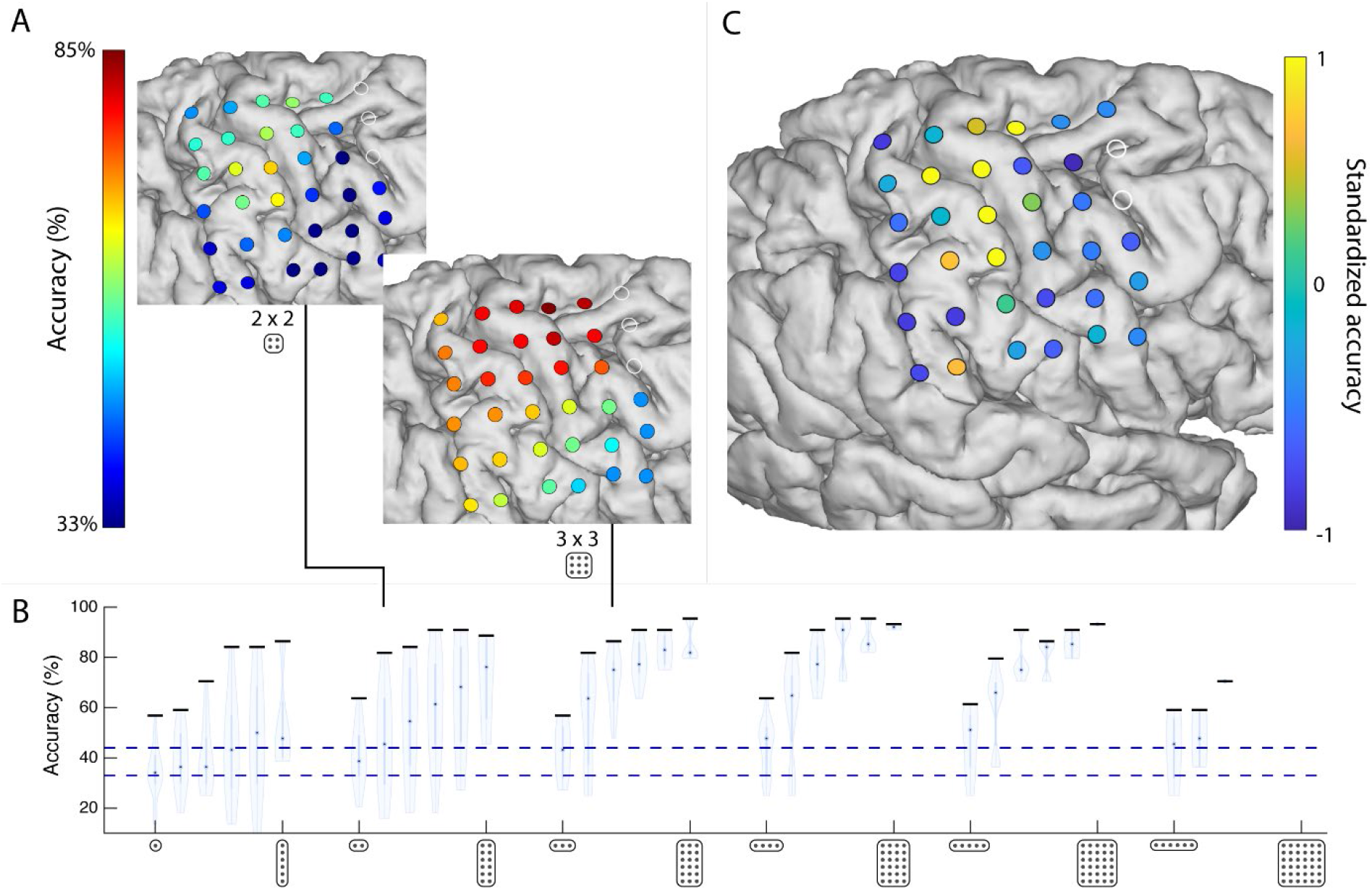
The spatial distribution of information. **(A)** Searchlight classification maps of searchlight with 2 × 2 (top) and 3 × 3 electrodes (bottom). **(B)** The classification results of the different searchlights, ranging from 1×6 to 6×6 grids in two directions (superior-inferior, anterior-posterior). Each violin plot represents the searchlight results with a specific grid configuration. The violin represents the distribution of the classification accuracies at the different searchlight positions within the grid, with a black horizontal bar to indicate the searchlight position that classified the highest. The lower dotted blue line shows the chance level at 33%, while the upper blue line indicates the threshold of 45% above which the decoding accuracy was significant. **(C)** Most informative electrodes, identified by a random search classification on 10.000 subsets of electrodes.

**Figure 6.**
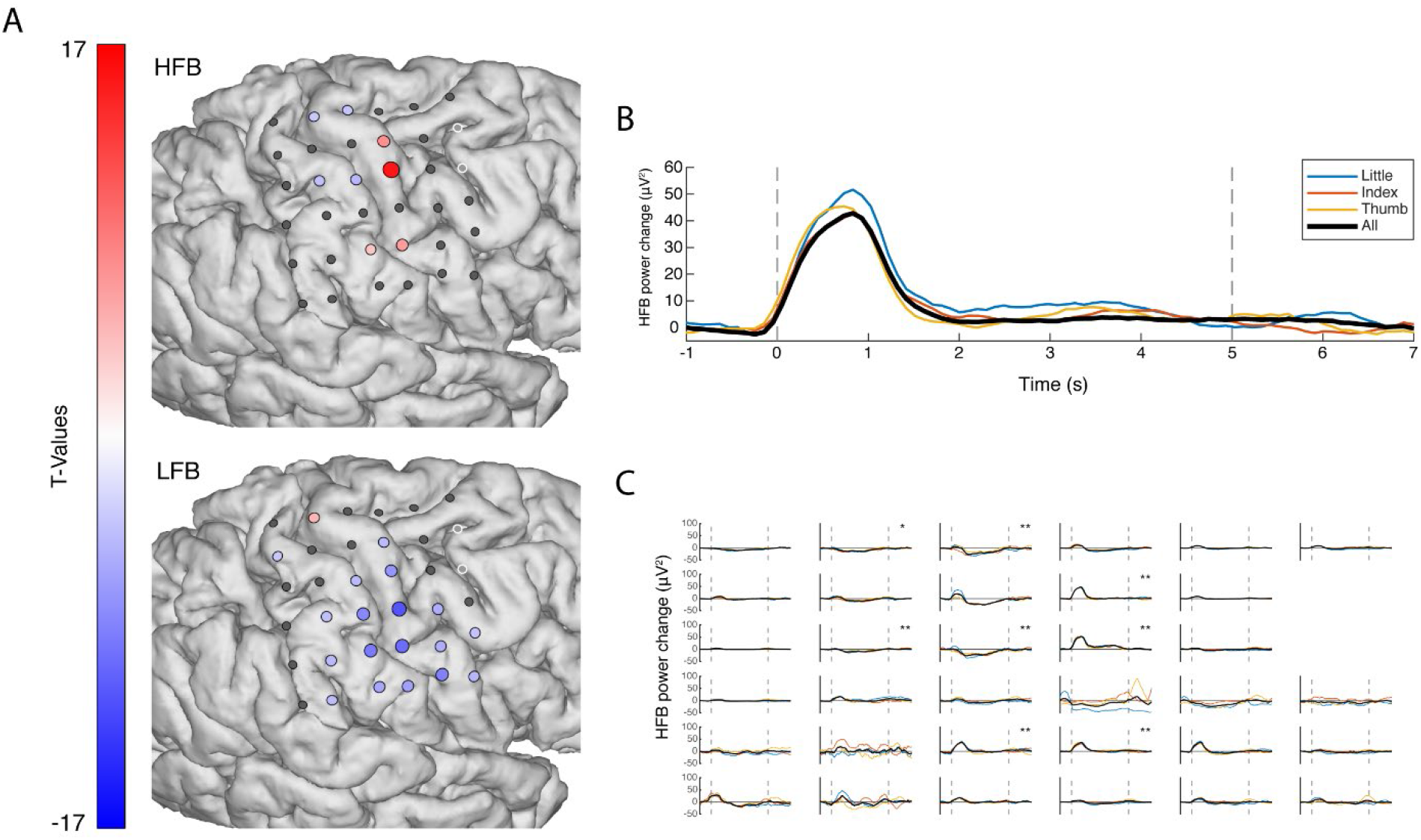
The, ipsilateral, electrophysiological response of executed movement versus rest. **(A)** The changes in HFB (top, 65-175Hz) and LFB power (bottom, 15-28Hz) for each grid electrode. Electrodes with a red or blue color showed a significant change in band-power, whereas electrodes with an insignificant change in band-power are shown in grey; the two excluded electrodes are shown in white. **(B)** HFB power changes over time averaged across those electrodes that showed a significant positive increase. The black line represents all fingers, whereas the colored lines represent individual fingers. The two vertical dotted lines indicated the cue on- and offset. **(C)** The HFB power changes over time, each graph represents one electrode.

To test the spatial extent of finger movement information, searchlight analyses with different types of grid configurations were performed (Figure 5A and 5B). Grid configurations with one or two electrodes tended to perform more poorly (~60%) compared to grids that include at least 3 electrodes (> 60%). For electrodes strips (i.e. grids that have multiple electrodes only in one dimension), the orientation of the grid becomes important. Grids with 3 to 6 electrodes that are placed along the superior-inferior axis perform much better at > 70% than grids are oriented on the anterior-posterior axis (at ~60%). This coincides with the topographical organization of the different fingers on the cortex, which is more superior-inferior oriented than anterior-posterior (Dechent and Frahm 2003; Miller et al. 2009; Siero et al. 2014; Schellekens et al. 2018; Huber et al. 2020). In accordance with the most informative area in Figure 5C, grids perform best when 2-3 electrodes wide and at least 2-3 electrodes high, in order to cover enough area of the brain to include the most informative electrodes. Given the inter-electrode distance of 10mm, the minimum required grid size to obtain a good (>= 80%) classification would be around 13mm × 13mm (2×2 electrodes).

### Ipsilateral finger representations

In addition to attempted movement, the patient also performed runs of executed hand movements with the healthy, ipsilateral, hand. Frame-perfect video annotations were used to quantify the movement of the healthy hand. An average delay of 0.44s (std: .09) occurred between the cue onset and actual start of the movement. The patient moved his hand for about 3.69s (std: 0.53) on average over all trials, with about 3-5 flexions of the finger per trial.

Only a few channels on the hand-area showed a strong ipsilateral increase in HFB power during executed movement, whereas some other channels showed a smaller, yet significant, ipsilateral decrease in HFB power. An ipsilateral distributed decrease in LFB was found, but was less spread out over the cortex compared to attempted movement. The HFB power for executed movement showed a transient response with a temporal peak between 0.5s and 1s, in line with the behavioral start of the hand movement and slightly earlier than the HFB peak in attempted movement at 1s.

Classification was performed on the ipsilateral response to the movement of the healthy hand. Based on the HFB power of all electrodes, finger movements could be classified with an accuracy of 56%.

## Discussion

In order to understand whether motor physiology is preserved after deafferentation, we investigated the contralateral electrophysiological responses of attempted finger movements in a patient with an upper-arm amputation. With attempted movement, a spatially-focal increase was found in broadband high-frequency ranges (65-175Hz) over the hand-area of the primary sensorimotor cortex. A spatially distributed decrease was found in the lower frequency bands (15-28Hz). Such an electrophysiological response is similar to that of executed finger and hand movements (Miller, 2009; Siero, 2014; Crone, 1998) suggesting that the motor physiology of the hand is retained.

Studies that investigated the motor electrophysiology in patients with locked-in-syndrome (i.e. ALS, PLS or brainstem-stroke) using EEG, MEG or permanent ECoG implants found similar results. In these patients, a robust HFB response was retained (Freudenburg et al. 2019). Whether and/or how the LFB response was affected varied between studies. Some studies observed robust low frequency power decreases in patients with ALS and/or PLS (Bai et al. 2010; Riva et al. 2012; Proudfoot et al. 2017). Other studies reported reduced power decreases in ALS (Kasahara et al. 2012), or more variability between patients with ALS, tetraplegia and brainstem stroke, with only some patients showing robust low frequency power decreases (Höhne et al. 2014; Freudenburg et al. 2019). These studies suggest that whether low frequency power decreases are retained depends on the disease (progression) and the influence of closed-loop feedback training. Here we have observed that there are strong and significant low frequency power decreases during attempted movement in a patient with an amputated arm.

Each individual finger resulted in a strong HFB response in the hand-region of the sensorimotor cortex. Although each of the fingers elicited a different HFB response pattern, no clear topographical representation of the fingers was found. Regardless, using support vector machine learning, we were able to decode the attempted movement of three individual fingers significantly at a classification accuracy of 93% (well above the 33% chance level). Which is similar to the decoding accuracies of executed movement of the fingers (Kubánek et al. 2009; Chestek et al. 2013). Attempted movements of the thumb could be decoded at 100% accuracy, however the index and little finger were less discriminable with sensitivity values of respectively 93% and 87%. Such decoding accuracies confirm that individual finger representations in the cortex are retained and can be distinguished in a patient with an amputated arm. Our results align with earlier fMRI research on patients with long term upper arm amputations (Bruurmijn et al. 2017), while other fMRI studies have shown displacement of the cortical activation into the deafferented motor and somatosensory areas during lip (Lotze et al. 2001), chin (Elbert et al. 1994) and face/shoulder movement (Dettmers et al. 2001). Two of these studies (Dettmers et al. 2001; Lotze et al. 2001) found that such displacements occur primarily in patients with phantom limb pain. It is possible that the cortical representations of other body parts invade the cortical regions of an amputated limb, which would warrant further investigation. However, studies in the visual cortex have shown only a limited ability for the primary cortex to reorganize (Smirnakis et al. 2005). Our research similarly demonstrates that the representation of the missing hand is at least largely retained and not replaced.

The HFB response of attempted movement was both transient and sustained, similar to what was found in research on continuous/repeated executed finger movements (Hermes, Siero, et al. 2012; Siero et al. 2013). For attempted movement, the HFB power peaked at ~1s after cue onset and returned gradually back to baseline during the remainder of the trial. Part of the latency between the cue onset and the peak of the cortical response can be explained by the lag between the interpretation of the cue and movement initiation, which in the executed movements of the patient already accounted for ~0.5s. Another factor that could have contributed to this ~1s latency may be related to the fact that motor imagery can be demanding in terms of mental fatigue and effort (Papadelis et al. 2007; Jacquet et al. 2020). In terms of decoding, both the transient and the sustained responses contained information about finger movements. The classifications of attempted finger movement over time confirmed that most information (i.e. the highest classification accuracies) was found around the peak of the response at 1s after cue presentation. The classification accuracies were more variable when including only the sustained response. However, larger time-windows (>2500ms) that included both the transient and sustained responses yielded higher classification accuracies than smaller windows (<2500ms), implying that the inclusion of (part of) the sustained response can contribute to the decodability.

In BCI applications, devices are often controlled with signals from the sensorimotor cortex using attempted movements (Collinger et al. 2013; Bouton et al. 2016; Vansteensel et al. 2016; Benabid et al. 2019). Our data showed that we could decode finger movements after deafferentation, suggesting that these signals can also be used to control a BCI. Understanding which signal properties allow for reliable decoding and BCI control is essential for these applications. Temporally, different parts of the electrophysiological signals can be included, but a tradeoff can occur between the speed of decoding and classification accuracy. A smaller window could allow for faster and more subsequent classifications, but could go at the expense of classification accuracy. Our data suggest that shorter (e.g. 1000ms) time windows may already provide a good accuracy (~80%) for decoding 3 fingers, while larger windows (e.g. 3000ms) will further improve accuracy (>90%). Patients may be able to use such short time windows, as one study in a patient with ALS already showed that movement versus rest can be decoded using a 1 sec window (Vansteensel et al. 2016). Understanding how well movement activity is retained after deafferentation may have implications for BCIs in patients with paralysis, as well as an amputated limb, as BCIs may reduce phantom limb pain (Yanagisawa et al. 2020).

Attempted movement after loss of function is different from movement imagery, and this distinction is of particular importance for implanted BCIs. It has been debated whether motor imagery representation overlaps with overt movement in brain surface recordings (Miller et al. 2010; Hermes et al. 2011), and whether motor imagery is a good approximation of attempted movement after limb loss or paralysis. One would intuitively expect that, in the case of a lost limb, the native map of representation would either be retained, or generally degrade. This patient’s map shows that somatotopic distinction is retained several years after limb loss. Some types of motor imagery in healthy individuals may thus not be a good general approximation or motor representations after limb loss or paralysis for implanted BCIs.

An additional point of importance for BCIs is the electrode grid design and extent of cortical coverage, which can have a strong influence on BCI performance (Vansteensel et al. 2016; Van Den Boom et al., 2021). Our results show that most information about attempted movement is located on the hand-region of the primary motor and sensory cortex. In terms of cortical coverage, considering an inter-electrode distance of 10mm, a good (>80%) classification accuracy can already be achieved with as little as 2×2 electrodes (13mm x 13mm) placed over the primary sensorimotor cortex. More electrodes could provide up to ~90% classification accuracy.

Finger movement activity on the ipsilateral cortex of the intact hand could be decoded, but less accurately compared to decoding the contralateral attempted finger movements. Only a few channels showed power increases during ipsilateral movements, while some electrodes also showed significant high frequency power decreases. Whether previous ECoG studies show similar ipsilateral high frequency power decreases during executed hand movements is less clear (Zanos et al. 2008). However, ipsilateral decreases in sensorimotor activity during hand movement in healthy subjects have been observed in the fMRI BOLD signal (Hlushchuk and Hari 2006; Diedrichsen et al. 2013). TMS studies similarly show evidence for contralateral inhibition (Talelli, Ewas, et al. 2008; Talelli, Waddingham, et al. 2008). The ipsilateral decreases in high frequency power we observed with ECoG may thus potentially be related to inhibition resulting from activity of the contralateral hemisphere, or to some reorganization of function after the injury (as has been seen with patients after perinatal hemispheric stroke; Miller et al. 2011)

### Conclusion

The electrophysiology of attempted hand movement is preserved in the sensorimotor cortex after deafferentation of an amputated hand, with a typical focal increase of HFB power over the hand region and a more distributed decrease in LFB. Attempted finger movements provided a transient HFB peak around 1s after cue onset, followed by a sustained HFB response. Classification analyses confirm that most decodable information on the finger movement can be found around this peak. Furthermore, HFB power can be used to decode finger movements with high (>90%) accuracy. Optimal decoding could be achieved based on the first 1-3s of the signal and would only require 13-13mm of cortical coverage. Our results demonstrate that the sensorimotor electrophysiology remains largely intact after long term (3 years and 11 months) amputations and therefore remains a viable region for BCIs that use the decoding of hand-gestures for control.

## Acknowledgements

We thank the patient and staff for their time and effort, and we would like to acknowledge Cindy Nelson and Karla Crockett for technical support.

This work was supported by NIH-NIMH CRCNS R01MH122258-01 (DH, KJM, MvdB), NIH-NIMH R01MH111417 (NFR) and the NIH-NCATS CTSA KL2 TR002379 (KJM). KJM is also supported by the Van Wagenen Foundation and by the Brain Research Foundation, with a Fay/Frank Seed Grant. GW & NMG are supported by NIH R01-NS092882 & R01-NS112144. NMG is also supported by the American Epilepsy Society Research & Training Fellowship for Clinicians. NFR was also supported by the European Research Council under award ERC-ADG 320708.

Manuscript contents are solely the responsibility of the authors and do not necessarily represent the official views of the NIH. None of the funding sources had any role in study design; in the collection, analysis and interpretation of data; in the writing of the report; and in the decision to submit the paper for publication.

## Supplementary material

**Supplementary Figure 1.**
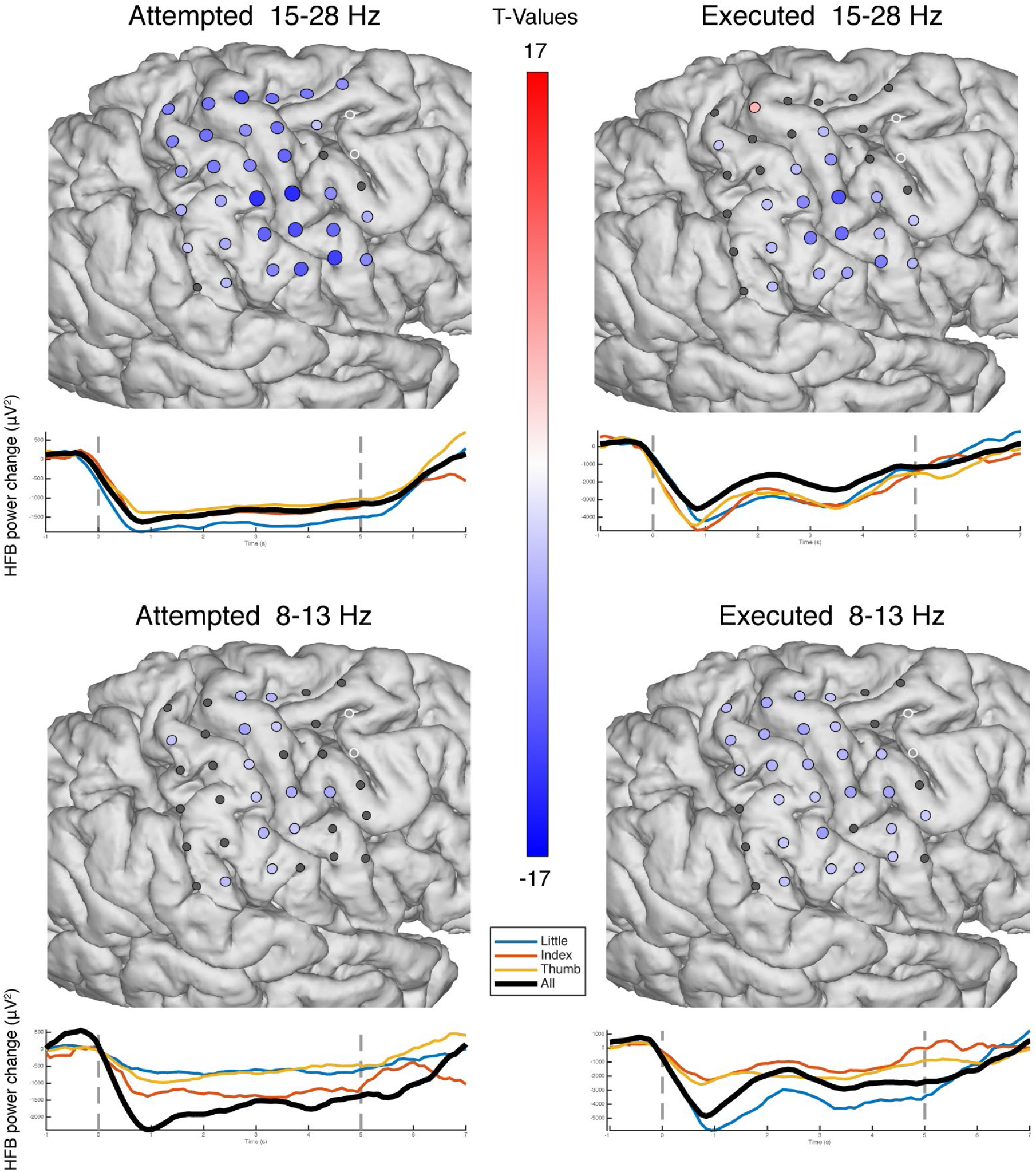
The electrophysiological responses in the spatial and time domain for both the 15-28Hz and 8-13Hz low-frequency bands during attempted movement (contralateral) and executed movement (ipsilateral). Electrodes with a red or blue color showed a significant change in band-power power, whereas electrodes with an insignificant change are shown in grey; the two excluded electrodes are shown in white. The time traces show the LFB power changes averaged across those electrodes that showed a significant decrease. The black line represents all fingers, whereas the colored lines represent individual fingers. The two vertical dotted lines indicated the cue on- and offset.

## Notes

https://osf.io/vmxdn/

### Competing Interest Statement

The authors have declared no competing interest.

